# β-catenin–independent regulation by TCF7L2 underlies isoform redundancy during embryonic thalamic development

**DOI:** 10.64898/2025.12.27.696670

**Authors:** Michael O. Gabriel, Joanna Bem, Marcin A. Lipiec, Abhishek Agarwal, Ewa Liszewska, Suelen Baggio, Huihui Qi, Dariusz Plewczynski, Marta B. Wiśniewska

## Abstract

Alternative promoter usage generates multiple transcription factor isoforms during brain development, yet their functional significance remains poorly defined. One such example is TCF7L2, a transcription factor critical for the development of the thalamus and recurrently affected by de novo mutations in autism spectrum disorder. TCF7L2 exists in two isoforms driven from different promoters: the long isoform (L-TCF7L2) with the β-catenin-binding domain, and the shorter isoform (S-TCF7L2), lacking this domain and classically considered a dominant-negative regulator of the L isoform. We investigated the role of TCF7L2 isoforms in thalamic development using total and isoform-specific knockout strategies. Integrated phenotypic and transcriptomic analyses revealed functional redundancy of TCF7L2 isoforms during embryogenesis. β-catenin subcellular localization and chromatin occupancy uncovered a developmental switch in TCF7L2 activity, from a β-catenin–independent and isoform-redundant mode in the embryonic thalamus to a β-catenin–dependent program postnatally. More broadly, these findings point to distinct embryonic and postnatal regulatory strategies, with alternative promoter usage potentially supporting robust availability of regulatory proteins during brain development.

## Introduction

Alternative promoter usage is widespread during brain development and changes markedly during the perinatal period (Pal et al., 2011; Tan & Sun, 2025; P. Zhang et al., 2017), when neuronal gene expression programs are extensively remodeled (Stroud et al., 2020). However, the functional contribution of alternative promoter usage to neural development remains poorly defined. One such example is TCF7L2.

TCF7L2 is a transcription factor and an effector of the canonical Wnt/β-catenin signaling pathway. It plays a key role in regulating gene expression during neuronal differentiation, as well as in the morphological and functional maturation of the developing brain. Recently, de novo mutations in *TCF7L2* have been linked to a rare neurodevelopmental disorder characterized by speech, motor, and intellectual disabilities, as well as features of autism spectrum disorder (Dias et al., 2021), highlighting the gene’s critical role in brain development.

TCF7L2 has been implicated in multiple aspects of brain development. It exhibits high and stable neuronal expression in the diencephalon and dorsal midbrain, in contrast to its transient expression in cortical progenitors and glial lineage cells (Chodelkova, Masek, Korinek, Kozmik, & Machon, 2018; Szewczyk et al., 2023). In these regions, TCF7L2 plays a critical role in establishing regional identity, connectivity and electrophysiological properties (Beretta, Dross, Bankhead, & Carl, 2013; Hüsken et al., 2014; Lee et al., 2017; Lipiec et al., 2020; Qi et al., 2023; Tran et al., 2020). Beyond this, TCF7L2 also regulates the expansion of the optic tectum in zebrafish (Brożko et al., 2022; Shimizu, Kawakami, & Ishitani, 2012), the neocortex in mice (Chodelkova et al., 2018), and the maturation of astrocytes and oligodendrocytes (Szewczyk et al., 2023; Zhao et al., 2016).

The complexity of TCF7L2’s action in the brain is further shaped by promoter usage which generates two isoform groups: full-length long isoforms (L-TCF7L2) containing the N-terminal β-catenin-binding domain, and short isoforms (S-TCF7L2) lacking this domain (Weise et al., 2010). L-TCF7L2 is broadly expressed in neuronal and non-neuronal tissues. In contrast, S-TCF7L2, transcribed from an internal promoter within intron 5, is specifically expressed in the diencephalon and dorsal midbrain.

S-TCF7L2 has been widely considered a dominant-negative isoform, acting as a competitive antagonist of the long isoform in Wnt signaling (Cadigan & Waterman, 2012). In this framework, S-TCF7L2 would be expected to counteract the actions of L-TCF7L2/β-catenin complex in brain regions where both isoforms are co-expressed. However, L-TCF7L2 does not always act through β-catenin-dependent mechanisms—for example, it regulates gene expression in differentiating oligodendrocytes independently of β-catenin(Iioka, Doerner, & Tamai, 2009; Ye et al., 2009). This raises the possibility that the S-TCF7L2 isoform may have functions in the thalamus beyond antagonizing Wnt/β-catenin signaling. Understanding this relationship is particularly important given that *TCF7L2* mutations identified in psychiatric patients are likely to affect both isoforms, as they are primarily located in exons shared by the L- and S-isoform transcripts (Coe et al., 2019; Dias et al., 2021; Iossifov et al., 2014; Satterstrom et al., 2020; Stessman et al., 2017; Szewczyk et al., 2023).

Here, we use genetically modified mouse models with total and isoform-specific knockouts of TCF7L2 to examine how individual TCF7L2 isoforms contribute to thalamic development. Our findings challenge the prevailing view of S-TCF7L2 as a dominant-negative isoform and reveal a developmental transition in TCF7L2 regulatory activity from β-catenin– independent mechanisms in the embryonic thalamus to β-catenin–dependent regulation in adulthood.

## Results

### Long and short TCF7L2 isoforms are present in neurons of the mouse thalamus throughout embryonic development

Previous research showed that both isoforms are expressed in the embryonic and adult thalamus, although their relative abundance changes postnatally (Nagalski et al., 2013). However, the dynamics of this change have not been analyzed. To determine the presence and distribution of TCF7L2 isoforms in the thalamus across developmental stages, we analyzed the spatiotemporal expression patterns in control mice using two antibodies: one detecting total expression of TCF7L2 (α-TCF7L2), which recognizes both isoforms, and another specific to the N-terminal β-catenin-binding domain of the long isoform (α-L-TCF7L2) (Fig. S1).

Immunofluorescence analysis revealed that the L-TCF7L2 isoform is present as early as embryonic day (E) 11.5 in the neurogenic thalamic primordium, with similar levels in the thalamic ventricular zone (VZ) and mantle zone (MZ). In contrast, total TCF7L2 levels were markedly higher in the MZ, indicating that the S-isoform is upregulated in postmitotic neurons. These findings show that both isoforms are present in the thalamus from the onset of postmitotic neurons (Fig. 1A).

**Figure 1.**
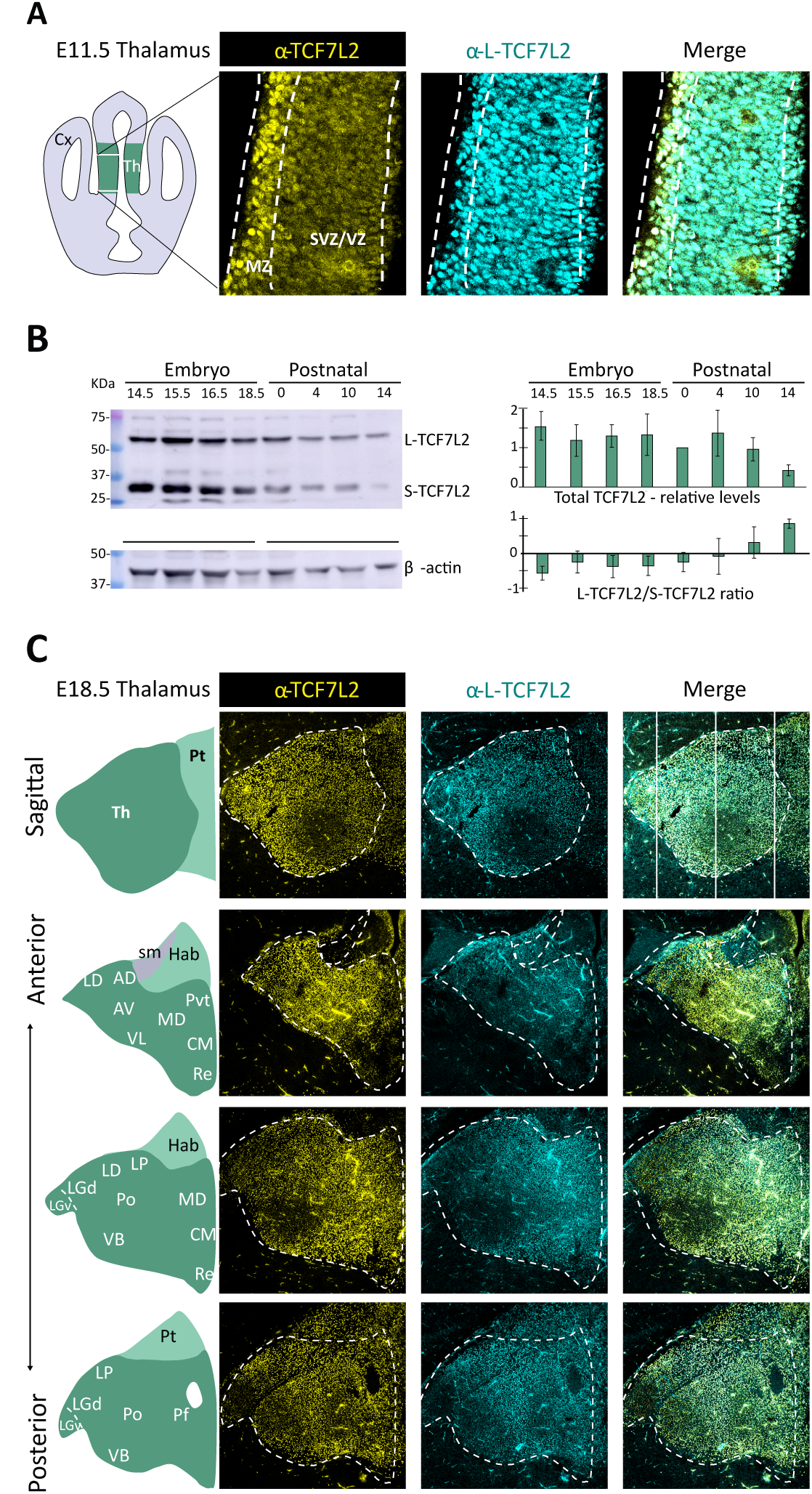
Both TCF7L2 isoforms are expressed in thalamic neurons throughout embryonic development. **[A]** immunostaining of TCF7L2 in thalamic neurons of E11.5 wild-type (WT) mice using antibodies that detect total TCF7L2 (both S- and L-TCF7L2) and L-TCF7L2. Higher signals of total TCF7L2 in the MZ versus SVZ/VZ suggest higher S-TCF7L2 in postmitotic thalamic neurons. **[B]** Representative immunoblot and quantification of TCF7L2 isoform levels across multiple prenatal and early postnatal stages of thalamic development in WT mice. Data presented as mean ± SD, n = 6. While both TCF7L2 isoforms are strongly expressed following thalamic neurogenesis, the relative abundance of the S-TCF7L2 declines markedly compared to the L-TCF7L2 at P14. **[C]** Immunostaining of TCF7L2 across thalamic nuclei areas mapped using the Allen mouse brain atlas (https://atlas.brain-map.org/) reveals broad yet spatially distinct expression patterns of TCF7L2 across major thalamic nuclei. AD - anterodorsal; AV -anteroventral; CM – centromedian; Hab - habenula; LD - lateral dorsal; lHb – lateral habenula; LGd - dorsal part of lateral geniculate; LGv – ventral part of Lateral geniculate; MD – mediodorsal; mHb - medial habenula; MZ - mantle zone; LP - lateral posterior; Pf - parafascicular; Po - posterior complex; Re - nucleus of reuniens; sm - stria medullaris; SVZ - subventricular zone; VB – Ventrobasal complex; VL – ventral lateral; LGv - ventral part of lateral geniculate; VM - ventral medial; VZ - ventricular zone.

To assess changes in TCF7L2 isoform composition over time, we quantified S- and L-TCF7L2 levels by Western blotting from E14.5 (the end of thalamic neurogenesis) to postnatal day (P) 14. TCF7L2 was expressed throughout embryonic and postnatal development. S-TCF7L2 was more prevalent during embryonic stages and declined sharply between P4 and P14, eventually falling below L-TCF7L2 levels (Fig. 1B).

As the thalamus comprises various nuclei with distinct molecular signatures (Govek et al., 2022; Nagalski et al., 2016), we examined the regional expression of TCF7L2 at E18.5 using the α-L-TCF7L2 and α-TCF7L2 antibodies. The L-TCF7L2 isoform was detected throughout the thalamus (Fig. 1C), with lower levels in the lateral and ventral regions, particularly in the anterior ventral nucleus (AV), lateral geniculate dorsal part (LGd), and the ventrobasal complex (VB) central area. Lower levels were also observed in the parafascicular nucleus (PF). Notably, L-TCF7L2 expression was lowest in the lateral geniculate ventral part (LGv) – a GABAergic region (Fig. 1C). Additionally, we observed lower L-TCF7L2 levels in the habenula. Total TCF7L2 exhibited a similar spatial distribution, indirectly suggesting comparable regional expression patterns of L-TCF7L2 and S-TCF7L2 across the thalamus and habenula.

In summary, the L-to-S TCF7L2 isoform ratio is higher in thalamic progenitors and the adult thalamus compared to the embryonic stage, where the S isoform is more abundant. This dual presence of the isoforms raises questions about the distinct functional roles of each isoform during the specific developmental window.

### Short TCF7L2 isoform is present in thalamic neurons of the L-TCF7L2-specific knockout mouse throughout embryonic development

The short isoform of TCF7L2 is traditionally associated with the inhibition of Wnt signaling, potentially protecting thalamic cells from excessive Wnt activity. However, it may also serve additional roles beyond inhibiting the long isoform. To explore these possibilities, we generated an L-TCF7L2-specific knockout (L-KO) in mice and compared its effects with those of the previously developed total TCF7L2 knockout (Lipiec et al., 2020).

L-KO was generated using CRISPR/Cas9 to delete a cytosine in exon 2, introducing a premature STOP codon (Fig. 2A). Sanger sequencing confirmed the mutation, and Western blotting showed the absence of the L-TCF7L2 isoform in thalamic lysates. At the same time, the S-TCF7L2 isoform was preserved (Fig. 2B-C). At E11.5, TCF7L2 immunofluorescent signal was detected throughout the thalamic primordium in L-KO embryos, with the highest intensity in the mantle zone (MZ) (Fig. 2D), consistent with the postmitotic induction of the short isoform. At E18.5, the immunofluorescence signal was present in thalamic regions, confirming the presence of the short TCF7L2 isoform throughout embryonic development (Fig. 2E).

**Figure 2.**
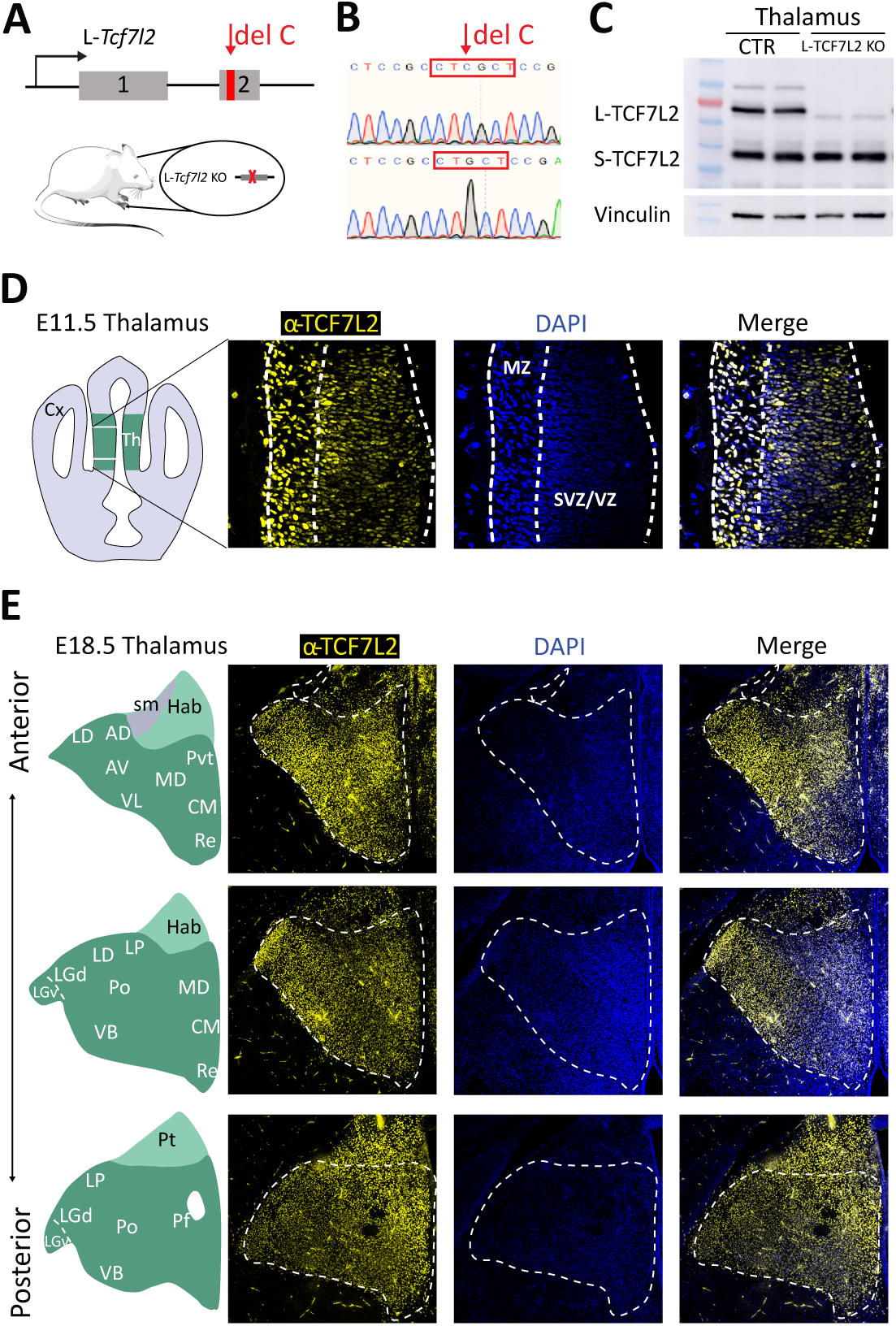
Validation of L-TCF7L2 deletion and localisation of S-TCF7L2 in the mouse thalamus. **[A]** Schematic of L-KO generation. **[B]** Chromatogram showing allele-specific deletion at the TCF7L2 locus. **[C]** Immunoblot showing that the 2569 antibody detects only the S-TCF7L2 isoform in L-KO mice. **[D]** Immunostaining revealed S-TCF7L2 is expressed in thalamic progenitors in the SVZ/VZ and increases in postmitotic neurons in the MZ at E11.5. **[E]** At E18.5, S-TCF7L2 is enriched in LGd/v, VPL, and VPM nuclei.

### Isoform-specific knockout of L-TCF7L2 results in a phenotype similar to the total knockout, but less severe

To assess the impact of each knockout strategy on thalamic development, we compared thalamic anatomy, thalamocortical axon guidance, molecular identities of thalamic cells and the formation of molecular boundaries in the diencephalon, which have been well characterised in t-KO embryos (Lipiec et al., 2020). We reasoned that loss of L-TCF7L2 would result in a phenotype similar to t-KO embryos if the short isoform primarily acts as a dominant-negative variant.

Brain sections from E18.5 control (CTR), total knockout (t-KO), and L-TCF7L2 isoform-specific knockout embryos were stained with Cresyl Violet to visualize diencephalic morphology (Fig. 3A). An antibody recognizing GAD67, a marker of GABAergic neurons, was used to delineate the anteroventral and posterior limits of the glutamatergic thalamus and to identify the GABAergic lateral geniculate ventral nucleus (LGv) (Fig. 3B). In L-KO embryos, the diencephalon exhibited mild elongation, accompanied by a reduced lateroventral aspect of the thalamus, with a clear histological boundary remaining between the thalamus and the habenula. In contrast, t-KO embryos showed pronounced horizontal thinning and vertical elongation of the diencephalon, leading to complete fusion of the thalamus and habenula, consistent with previous findings.

**Figure 3.**
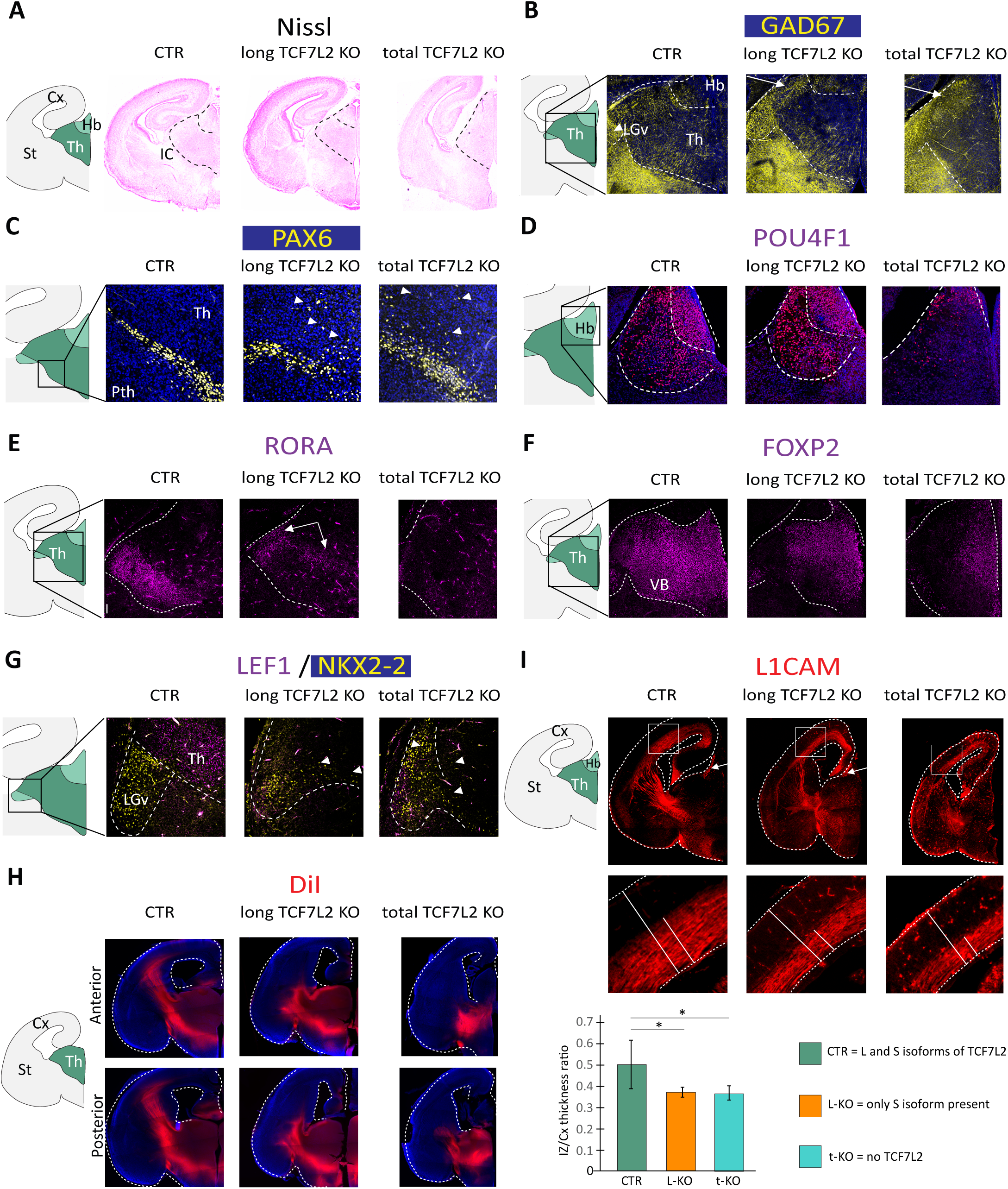
Loss of L-TCF7L2 recapitulates the global TCF7L2 KO phenotype with reduced severity at E18.5. **[A]** Nissl stain of mouse brain sections from Control, L-KO and t-KO models reveals markedly greater disruption of diencephalic morphology of t-KO compared to L-KO and control mice at. **[B]** GAD67 immunostaining used to delineate the boundaries of non-GABAergic thalamus from surrounding GABAergic areas in experimental models with altered diencephalic morphology. Immunodetection of **[C]** prethalamic marker, PAX6; and **[D]** habenular marker, POU4F1, revealed selective disruption of the ventral thalamic border, whereas t-KO mice exhibited pronounced alterations at both ventral and dorsal thalamic borders. Immunostaining for the thalamic regional and sub-regional markers **[E]** RORA; **[F]** FOXP2; **[G]** LEF1 and NKX2-2, reveals a marked decline in expression across both L-KO and t-KO mice compared to control. **[H]** DiI axon tracing revealed a severe disruption in thalamocortical projections in L-KO mice and more dramatically in the t-KO mice. **[I]** L1CAM immunostaining and IZ morphometry showed a significant thinning of the IZ relative to cortical thickness in both t-KO and L-KO mice compared to the control. Cx – cortex; Hb – habenula; IZ – intermediate zone of the cortex; LGd - dorsal part of lateral geniculate; LGv – ventral part of Lateral geniculate; Pth – prethalamus; St – striatum; Th – thalamus

To further delineate thalamic boundaries in L-KO embryos, we performed immunofluorescent staining of regional markers: PAX6 for the ventrally located prethalamus and POU4F1 for the dorsally located habenula (Fig. 3C-D). In both knockouts, prethalamic and thalamic cells intermingled at the ventral boundary, indicating that L-KO embryos exhibited a similar disruption of the dorsal thalamic border as observed in t-KO embryos. However, in L-KO embryos, the dorsal boundary and habenular identity were maintained, unlike in t-KO embryos.

To assess the impact of L-KO on the subregional molecular identities of the E18.5 thalamus, we examined markers of the glutamatergic thalamus: RORA, FOXP2 and LEF1; and the GABA-ergic thalamic (LGv) marker NKX2_2. At this developmental stage, RORA is expressed in the ventrobasal complex; this expression was partially preserved in the lateral and dorsal regions of the VB in L-KO mice but was lost in t-KO mice (Fig. 3E). FOXP2 expression, normally broad in the thalamus, was significantly reduced in the lateral and ventral thalamic nuclei in both knockout models (Fig. 3F), and LEF1 expression was absent in both L-KO and t-KO mice (Fig. 3G). While NKX2-2-positive neurons were present in the thalamus of control embryos and both knockout models (Fig. 3G), they were no longer clustered in the LGv region but instead spread along the lateral thalamic wall and dorsally in L-KO and t-KO mice. These results indicate that the L-TCF7L2 isoform plays a crucial role in specifying thalamic subregional identities, as its loss recapitulates the molecular phenotype observed in the complete knockout, albeit with reduced severity.

Next, we analyzed efferent fibers projecting from the thalamus to the cortex (thalamocortical axons, TCAs). To visualize these tracts, we traced them by inserting lipophilic DiI crystals into the thalamus. In L-KO embryos, TCAs, which should have reached the cortex by E18.5, failed to extend beyond the subpallium, indicating a partial impairment of thalamocortical projections (Fig. 3H). In contrast, in t-KO embryos, TCAs were virtually absent from the subpallium, reflecting a more severe defect. Consistent with the DiI tracing, immunostaining of forebrain sections with an α-L1CAM antibody revealed that fewer TCAs exited the thalamus and entered the ventral telencephalon in L-KO embryos compared with controls, whereas only sparse fibres were detected in the subpallium of t-KO embryos (Fig. 3I). Yet, both knockout models exhibited a similar thinning of the cortical intermediate zone, suggesting that TCAs also failed to reach the cortex in L-KO embryos.

In summary, our comparative phenotypic analysis of L-KO and t-KO embryos reveals that although both knockout strategies result in significant disruptions of thalamic development, the L-TCF7L2 isoform knockout produced a similar but milder phenotype compared to the total knockout. These findings suggest that the S-TCF7L2 isoform contributes to thalamic development in a way that extends beyond simply inhibiting L-TCF7L2, leading us to investigate the individual effect of each isoform on gene expression

### L- and S-TCF7L2 isoforms play redundant roles in transcription regulation in the embryonic thalamus

To determine whether the L- and S-TCF7L2 isoforms have opposing, redundant, or distinct roles in transcription regulation during thalamic development, we analyzed the effects of total and isoform-specific TCF7L2 knockouts (L-KO and S-KO) on the thalamic transcriptome at E18.5 using bulk RNA sequencing. S-KO embryos were generated by deleting the transcription start sites of S-TCF7L2 transcripts within intron 5 of the *Tcf7l2* gene, as previously described (Qi et al., 2023). If the isoforms have opposing functions, with S-TCF7L2 acting as an inhibitor of L-TCF7L2, we would expect differentially expressed genes (DEGs) to change in opposite directions in S-KO and L-KO. Conversely, minimal or no changes in either knockout would suggest functional redundancy. A lack of overlap between the sets of genes disrupted by the short and long isoform knockouts would suggest that each isoform plays a complementary and independent role in thalamic development.

Differentially expressed genes (DEGs) were identified by comparing thalamic transcriptomes between knockout and control groups, using a q-value threshold of < 0.05 and log₂ fold change (|log₂FC|) ≥ 0.4. In total 297 DEGs were found in t-KO embryos, whereas L-KO and S-KO embryos displayed fewer DEGs: 79 and 37, respectively. Most L-KO and S-KO DEGs (65 and 22, respectively) overlapped with those observed in t-KO embryos (Fig. 4A).

**Figure 4.**
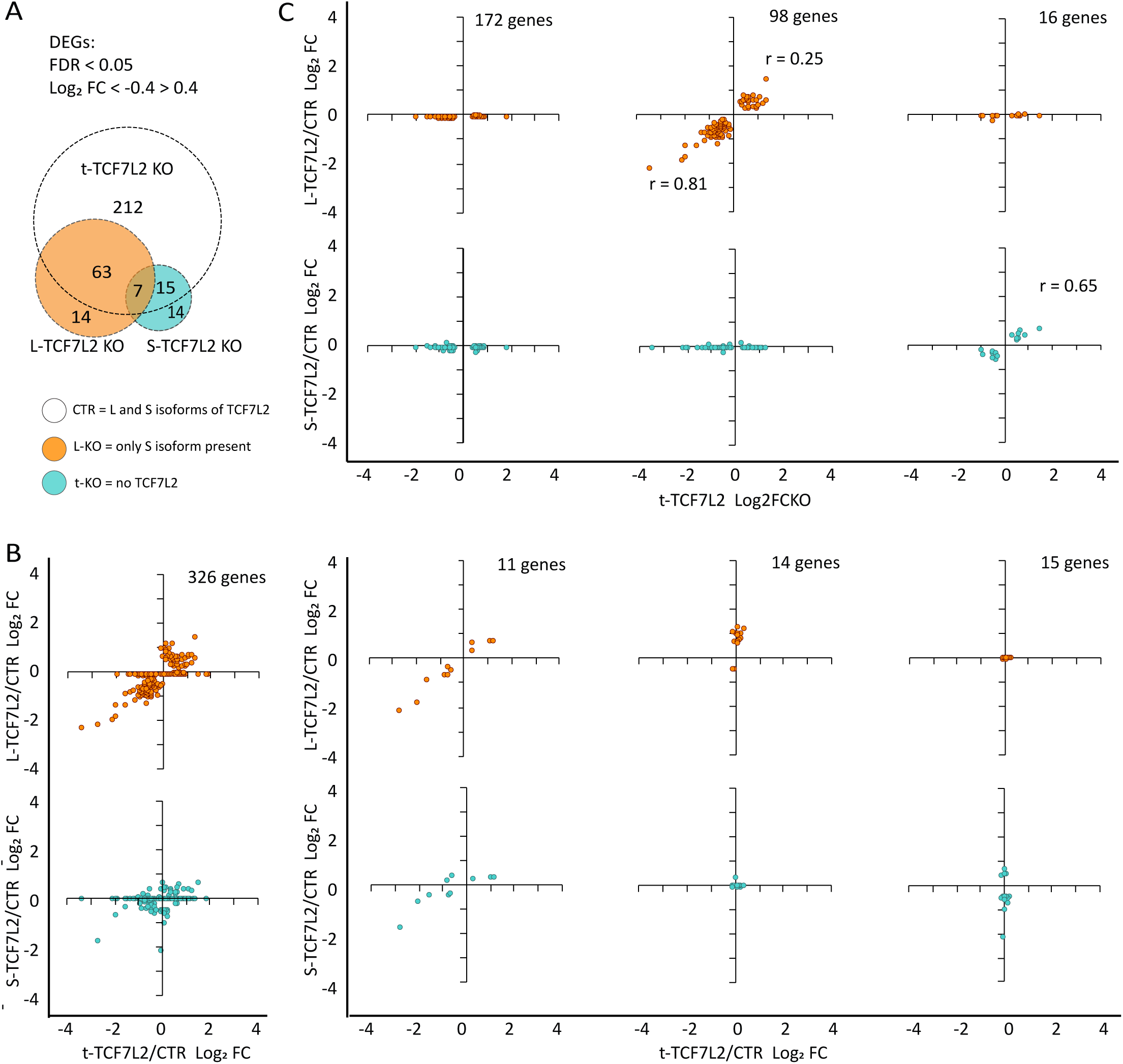
Transcriptomic changes following total and isoform-specific loss of TCF7L2 in the thalamus. **[A]** Venn diagram illustrating the intersection of differentially expressed genes (DEGs) with Log2 fold change ≤ −0.4 and ≥ 0.4 across t-KO, L-KO and S-KO mouse models at E18.5. **[B]** Scatter plots depicting the full set of 326 DEGs with significant expression changes (Log2 fold change ≤ −0.4 and ≥ 0.4) in at least one of the three knockout models. **[C]** Subset of DEGs altered in the t-KO mice but not in the L-KO or S-KO mice (suggesting redundant regulation by both TCF7L2 isoforms), or altered in the t-KO mice and only one of L-KO or S-KO (indicating isoform-preferential regulation). **[D]** Rare subset of genes altered exclusively in L-KO or S-KO models or in all three models, potentially representing isoform-independent changes or experimental artefacts due to their low occurrence.

To further explore the effects of single isoform knockouts, we compared the direction of Log_2_ FC for all significant DEGs across all three models (326 genes Fig. 4B, STable 1). This analysis revealed several distinct gene populations with different behaviours across the comparisons (Fig. 4C). The largest population of DEGs (172) showed altered expression exclusively in the t-KO model but not in either isoform-specific knockout, indicating high redundancy between the L- and S-isoforms. The second-largest population of DEGs (98) exhibited changes in both L-KO and t-KO conditions, but not in S-KO embryos (Fig. 4C), suggesting a more substantial influence of the L-isoform on TCF7L2-mediated gene expression. A smaller group (16) showed altered expression in t-KO and S-KO conditions, and a few genes (11) were affected across all three knockout models. Most DEGs in all populations displayed lower FC values and failed to reach statistical significance in isoform-specific knockouts (Fig. 4C, Table S1), supporting the notion that the remaining isoform compensates for the loss of the other. Notably, all expression changes occurred in the same direction, arguing against the opposing role of the L- and S-isoforms.

In conclusion, our results support the concept of redundant gene expression regulation by the L- and S-TCF7L2 isoforms in the developing thalamus. Based on these findings, we next asked whether TCF7L2 mediates the canonical Wnt/β-catenin pathway in this context.

### TCF7L2-mediated gene expression regulation is independent of β-catenin during thalamic development

The observed redundancy in gene expression regulation by TCF7L2 isoforms suggests that their function may be independent of β-catenin. To investigate this possibility, we examined β-catenin nuclear localization—a hallmark of canonical Wnt signaling—in the thalamus at E18.5 and subsequent developmental stages using immunofluorescence. In addition to its nuclear role in gene expression regulation, β-catenin also anchors transmembrane cadherins to the cytoskeleton (van der Wal & van Amerongen, 2020), therefore we also evaluated its membranous localization.

While a strong β-catenin membrane signal was detected at all developmental stages, nuclear β-catenin was absent at E18.5 and P4 (Fig. 5A), indicating limited β-catenin availability for TCF7L2 in the embryonic thalamus. Between P14 and P60, β-catenin gradually appeared in TCF7L2-positive cell nuclei, coinciding with a relative decrease in S-TCF7L2 isoform levels (Fig.1B). These findings suggest an increasing reliance of TCF7L2 on β-catenin in the postnatal thalamus.

**Figure 5.**
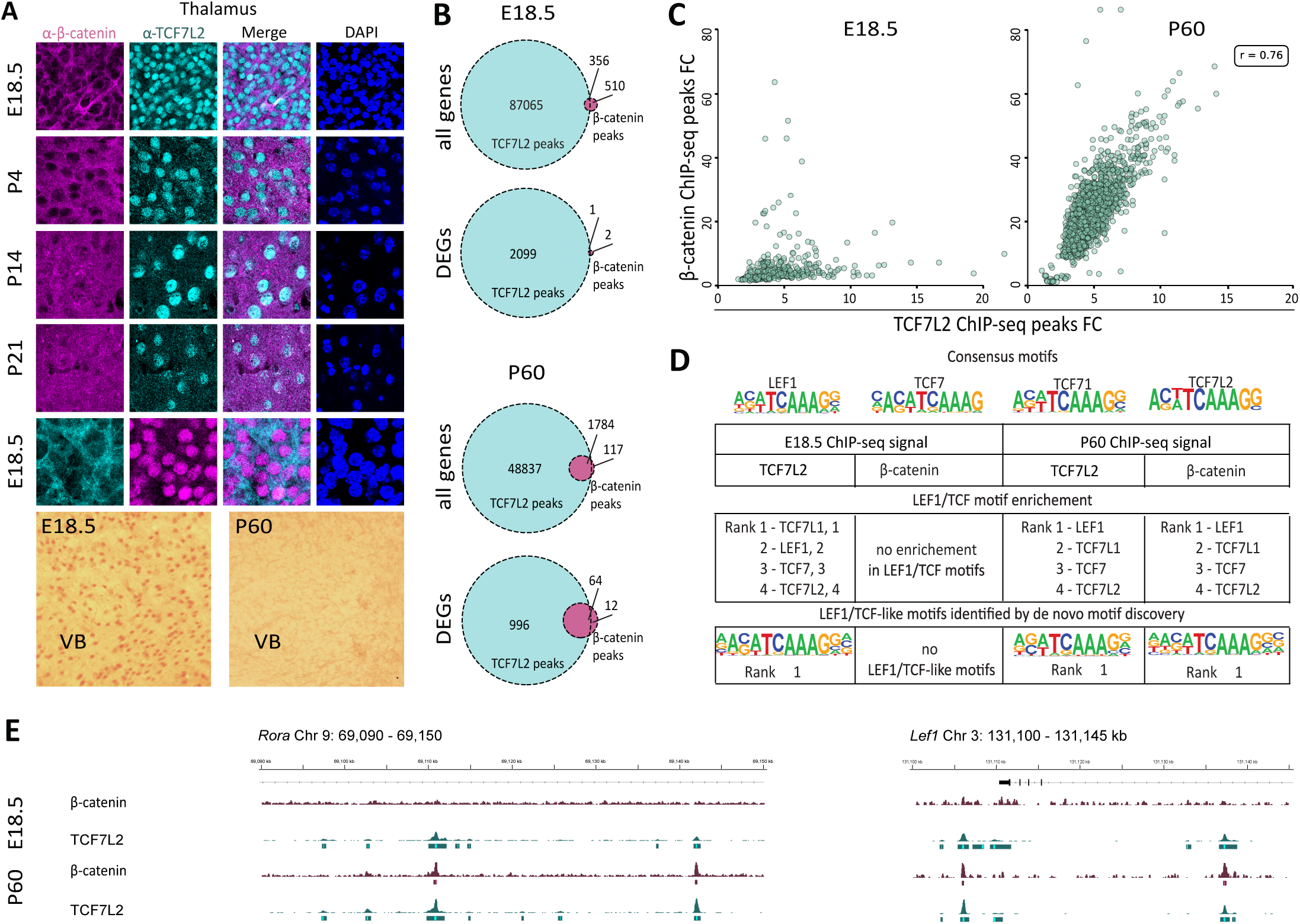
TCF7L2 regulates thalamic development independently of β-catenin. **[A]** Immunostaining showing the expression of TCF7L2 and β-catenin in the thalamus across prenatal, early postnatal (P4, P14) and adult (P60) stages. Signal of Nuclear β-catenin increases markedly at P14 and in adult (P60) compared to E18.5 and P4. **[B]** Venn diagram illustrating the intersection between CHIP-seq-identified gene targets that co-localise with TCF7L2 and β-catenin at E18.5 and P60. A significantly lower overlap at E18.5 compared to P60 suggests minimal interaction between TCF7L2 and β-catenin during embryonic development. **[C]** Correlation analysis between fold-change enrichment of TCF7L2 and β-catenin peaks revealed a strong correlation at P60 with no detectable concordance at E18.5 **[D]** Motif enrichments and de novo motif discovery analysis showed the absence of Lef1/TCF motif in β-catenin-bound chromatin at E18.5 in contrast to robust enrichment at P60, supporting a model in which TCF7L2 regulates gene expression independently of β-catenin during embryonic thalamic development. **[E]** Genome browser views of the Rora and Lef1 loci showing ChIP-seq signal for TCF7L2 and β-catenin at E18.5 and P60. Co-localization of β-catenin and TCF7L2 peaks is observed at P60, whereas β-catenin peaks are absent at E18.5.

To determine the cooperation between TCF7L2 and β-catenin in gene expression regulation in the thalamus, we performed ChIP-seq analyses using α-TCF7L2 and α-β-catenin antibodies. ChIP-seq data for P60 have been previously published (Lipiec et al., 2020). At E18.5, we identified 87,421 TCF7L2 peaks and 866 β-catenin peaks using a q-value threshold for signal fold-change enrichment > 0.05 (Fig. 5B). At P60, the number of TCF7L2 peaks was roughly half that at E18.5, 50,621, whereas β-catenin peaks doubled, with 1,901 detected using the same threshold. The generally lower number of β-catenin peaks compared to TCF7L2 likely reflects the indirect nature of β-catenin-chromatin binding, which reduces precipitation efficiency.

We next examined whether β-catenin colocalizes with TCF7L2 by overlapping their chromatin peaks. At E18.5, only 0.05% of TCF7L2 peaks overlapped with β-catenin peaks (Fig. 5B), and when the analysis was restricted to DEG regions, only one TCF7L2 peak overlapped with a β-catenin peak. No correlation was observed between the fold-change enrichment of TCF7L2 and β-catenin signals (Fig. 5C). These results suggested a weak to absent interaction between these two proteins in the embryonic thalamus.

At P60, in striking contrast to E18.5, nearly all β-catenin peaks overlapped with TCF7L2 peaks. (Fig. 5B). This indicated a robust interaction between β-catenin and TCF7L2 in the mature thalamus, further supported by a correlation in fold-change enrichment between TCF7L2 and β-catenin peaks (Fig. 5C).

To further investigate the involvement of β-catenin in TCF7L2-mediated gene regulation at E18.5 and P60, we examined the presence of TCF7L2 binding motifs within both TCF7L2 and β-catenin ChIP-seq peaks. Analysis of known motif enrichments and motif discovery in TCF7L2-bound chromatin revealed the strongest enrichment of LEF1/TCF motifs at both E18.5 and P60 (Fig. 5D), validating the specificity of TCF7L2 binding. In contrast, LEF1/TCF motifs were absent in β-catenin-bound chromatin at E18.5, but became the most enriched motifs at P60, as illustrated at the *Rora* and *Lef2* loci (Fig. 5E), indicating stage-specific engagement of β-catenin with LEF1/TCF-responsive elements. These results further indicate that β-catenin does not participate in TCF7L2-mediated gene expression regulation in the embryonic thalamus.

These findings reveal a shift in TCF7L2’s regulatory mechanisms, from β-catenin-independent during development to β-catenin-dependent in adulthood.

## Discussion

The results presented here refine the current understanding of how TCF7L2 operates during embryonic thalamic development. Although TCF7L2 is widely regarded as an effector of canonical Wnt/β-catenin signaling, our data reveal that the functional logic of TCF7L2 isoforms in the embryonic thalamus differs from the antagonistic model inferred from canonical Wnt signaling.

Both long and short TCF7L2 isoforms are abundant in the thalamus throughout embryogenesis, followed by postnatal downregulation and a shift toward long isoform predominance shown here and in previous papers (Lipiec et al., 2020; Nagalski et al., 2013). This developmental change coincides with a period marked by major shifts in gene expression profiles in neurons (Stroud et al., 2020), when promoter usage in the brain changes markedly between late embryogenesis and the early postnatal period (Pal et al., 2011; Tan & Sun, 2025; P. Zhang et al., 2017). TCF7L2 thus follows this general pattern.

During embryogenesis, when both TCF7L2 isoforms are co-expressed at high levels, their relative contributions to thalamic development can be directly interrogated. Functional analyses using total and long isoform-specific knockout revealed that loss of the long isoform only partially phenocopies the complete knockout. This graded phenotypic severity indicates that the short isoform partially compensates for the loss of the long isoform in multiple developmental processes. Notably, some aspects of diencephalic organization and patterning—intact habenular identity and preservation of the dorsal thalamic boundary—are fully preserved in the absence of the long isoform, indicating that the short isoform can fully substitute for long isoform function in some contexts. At the same time, similar disruption of the ventrolateral thalamic boundary between glutamatergic and GABAergic regions in both total and long isoform knockouts indicates that other aspects of thalamic development are more susceptible to loss of the long isoform. These observations underscore that while redundancy represents the dominant mode of action of TCF7L2 isoforms in the embryonic thalamus, it is not absolute equivalence.

In the canonical Wnt signaling framework, TCF7L2 activity is mediated through β-catenin, and short isoforms lacking the β-catenin–binding domain are therefore expected to function as competitive inhibitors of the full-length protein. Under this model, knockout of the long isoform would be predicted to produce effects opposite to those caused by loss of the short isoform. However, the partial phenotypic rescue observed in long isoform knockouts is inconsistent with this expectation. Moreover, contrary to this prevailing view, single-isoform (long or short) knockouts did not produce gene-expression changes in opposite directions, thereby arguing against a model of competitive inhibition. Instead, dysregulated genes showed expression changes in the same direction across all knockout models, with a smaller magnitude in isoform-specific knockouts than in the total knockout. Notably, loss of the short isoform affected very few genes, whereas loss of the long isoform disrupted a larger—but still limited—subset of the transcriptional changes observed in the total knockout, suggesting a greater contribution of the long isoform. This pattern is consistent with functional redundancy and dosage-dependent regulation rather than antagonistic control between isoforms, suggesting that alternative TCF7L2 isoforms contribute to regulatory robustness during thalamic development.

The lack of antagonism between TCF7L2 isoforms is readily explained by the virtual absence of nuclear β-catenin in the embryonic thalamus. Under these conditions, TCF7L2-mediated transcription proceeds through β-catenin–independent mechanisms. This regulatory logic changes postnatally and in the adult thalamus, where nuclear β-catenin is abundant and extensively co-occupies chromatin with TCF7L2. Notably, however, even at this stage TCF7L2–β-catenin cooperation in the thalamus does not fully conform to the classical Wnt paradigm. As previously shown, β-catenin accumulates in the nuclei of thalamic neurons even when Wnt signaling is inhibited, and this nuclear localization depends on TCF7L2 itself, pointing to a non-canonical mode of β-catenin recruitment (Misztal, Wisniewska, Ambrozkiewicz, Nagalski, & Kuznicki, 2011). Although TCF7L2 is classically viewed as an effector of Wnt/β-catenin signaling, our findings suggest that TCF7L2 may act β-catenin-independently during brain development. This idea is further supported by previous studies showing that TCF7L2 can regulate gene expression independently of β-catenin through interactions with alternative transcriptional co-regulators during oligodendrocyte differentiation (Zhao et al., 2016).

Our findings suggest a broader role for alternative promoter usage during brain development. Rather than primarily diversifying transcriptional outputs, alternative promoters may in some contexts serve to ensure sufficiently high and sustained levels of key regulatory proteins during periods of rapid developmental change. In the case of TCF7L2, alternative promoter usage generates isoforms that differ structurally, yet act redundantly during embryonic thalamic development. Such a mechanism may be particularly relevant during embryogenesis, when developmental programs must be executed reliably.

This regulatory model may be relevant for neurodevelopmental disorders. De novo mutations in *TCF7L2* identified in autism spectrum disorder affect exons shared by both isoforms and are frequently predicted to generate truncated proteins in a heterozygous state (Coe et al., 2019; Dias et al., 2021; Iossifov et al., 2014; Satterstrom et al., 2020; Stessman et al., 2017; Szewczyk et al., 2023), supporting a dosage-sensitive rather than isoform-specific disease mechanism. In line with this interpretation, our results show that embryonic thalamic development depends on the combined activity of both *TCF7L2* isoforms, consistent with a model in which reduced overall *TCF7L2* levels may contribute to disease-associated phenotypes.

Together, these findings suggest that alternative promoter usage can support robustness of transcriptional control during brain development, providing a regulatory framework that may be particularly sensitive to dosage perturbations associated with neurodevelopmental disorders.

## Materials and Methods

### Animal models

Animal maintenance and experimental protocols adhered to the standards of the Polish government and European community on animal experimentation and were closely supervised by the Animal Welfare Advisory Board of the Centre of New Technologies, Warsaw. Mice were bred under standard conditions with 12-h/12-h light/dark cycle and unrestricted access to food and water. The protocol for generating the TCF7L2 ^-^/^-^ mice (t-KO) which carries the *tm1a(KOMP)Wtsi* allele with LacZ cassette upstream of exon 6 of the *Tcf7l2* gene and the S-TF7L2 ^-^/^-^ mice (S-KO) carrying a mutation in the transcription start sites of the S-TCF7L2 isoform have been previously described (Lipiec et al., 2020; Qi et al., 2023). To create the L-TCF7L2 ^-^/^-^ mice (L-KO), a single nucleotide deletion was introduced to the second exon of the *Tcf7l2* gene using the CRISPR/Cas9 approach, producing an early stop codon in the gene’s reading frame.

### Tissue preparation and immunofluorescence

Embryos from pregnant E18.5 mice were collected and the brain tissues were removed and placed in 4% PFA for a whole day at 4°C to be fixed. The fixed brains were then cryoprotected for 72 hours at 4°C by transferring them to 30% sucrose in PBS. Tissues were imbedded in a solution of 30% sucrose and 30% gelatin in PBS (Ferran et al., 2015) and left to harden in order to prepare them for cryosectioning. Then, the embedded tissues were frozen in cold isopentane at about −40°C and then transferred to a freezer at −80°C for storage. Using a cryostat (Leica CM1860, Germany), tissue blocks were sectioned at 20 µm for embryonic tissues. Before staining, sections were placed on 25 x 75 x 1.0 mm Superfrost® Plus glass slides, which were kept at −20°C.

Brain tissue cryosections on glass slides were taken out of a storage freezer and allowed to defrost and dry at ambient temperature. The sections were submerged in warm (37–40°C) 1X PBS for 5 minutes with gentle rocking to wash off the gelatin, and then again for 10 minutes each in PBS-T. Sections were submerged in heated (70°C) citrate buffer to expose the antigen at room temperature for 40–45 minutes. Filtered 5% normal donkey serum (NDS) containing 0.3 M glycine in 1X PBS was used to block tissue slices. Thereafter, sections were treated in a cocktail of primary antibodies overnight at 4°C. These antibodies target all TCF7L2 isoforms (Cell Signalling Technology, 2569S, rabbit mAb, 1:1000) or only L-TCF7L2 (TCF-4, Merck Millipore, 05-511, mouse, 1:1000), PAX6 (BioLegend, 901301, rabbit, 1:200), RORA (Cell Signalling Technology, 34639S, rabbit, 1:1000), L1CAM (Merck Millipore, MAB5272, Rat, 1:500), NKX2-2 (DSHB, 74.5A5, mouse, 1:50), FOXP2 (Cell Signalling Technology, 5337S, rabbit, 1:250), POU4F1 (1:500; (Fedtsova & Turner, 1995), LEF1 (Cell Signalling Technology, C18A7, rabbit, 1:300), β-catenin (BD Transduction Lab, 610154, mouse, 1:200). Next, sections were rinsed three times in PBS-T for ten minutes each, then, sections were incubated for about an hour at room temperature in secondary antibody diluted in 1% NDS (1:500). The secondary antibodies includes Alexa Fluor^TM^ 594 or 488 (A-11058, A-21202, A-21203, A-21206, A-21207, A21209) from ThermoFisher Scientific. Subsequently, the sections underwent three 10-minute PBS-T washes before being cover-slipped using Fluoromount-G with DAPI (00-4959-52, Thermo Fisher Scientific). The confocal microscope was used to capture the images.

### DiI Axon Tracing

DiI axon tracing protocol followed a previously used method (Lipiec et al., 2020). This involves fixing E18.5 mouse embryonic brains in 4% PFA for 24 hours, followed by dissection into two hemispheres. Each thalamus received 1,1’-Dioctadecyl-3,3,3’,3’ Tetramethylindocarbocyanine Perchlorate (DiI) crystals (D3911, Thermo Fisher Scientific). Then, the brain tissues with the DiI crystals were incubated in 4% PFA for 21 days at 37°C with light protection. Thereafter, the brains were embedded in 5% low-melting-point agarose, and 100 µm-thick coronal sections of the brain were produced using a vibratome. The cutting of the brain tissues was performed at roughly 45 degrees to the horizontal.

### Western blotting

The thalamohabenular region of E18.5 mice was excised according to the method previously described by our laboratory (Lipiec et al., 2020). The brain samples were homogenised in RIPA buffer containing protease and phosphatase inhibitor and allowed to rest on ice for 5 mins before sonication. Thereafter, the lysate was centrifuged and the total protein in the supernatant was used for BCA Bradford (Bio-Rad) assay and quantified with a Biophotometer D30 (Hamburg, Germany). Twenty micrograms of protein from each sample were separated by SDS-PAGE and transferred to a nitrocellulose membrane. The membrane was blocked for one hour in 10% low-fat milk and the membrane was incubated overnight in primary antibody solution against TCF-4 (Merck Millipore, 05-511, mouse, 1:1000); TCF7L2 (Cell Signalling Technology, 2569S, rabbit mAb, 1:1000); Vinculin (Abcam, AB129002, rabbit, 1:3000) and β-actin (Sigma-Aldrich, A3854, mouse, 1:50,000). The membrane was washed three times for 5 minutes each in TBS-T and incubated for two hours in secondary antibody: anti-Mouse IgG – Peroxidase (Sigma-Aldrich, A9044, rabbit 1:10,000) and Anti-Rabbit IgG – Peroxidase (Sigma-Aldrich, A0545, goat 1:10,000). The membrane was developed in ECL substrate, and the protein bands were visualised with Amersham imager 600 (GE Healthcare Bio-Sciences AB). Relative protein levels were quantified by densitometric analysis of background-subtracted protein band volumes using Bio-Rad Image Lab software (version 6.0.1).

### RNA isolation and sequencing

Total RNA from E18.5 mouse thalamohabenular region was extracted using QIAzol solution (Qiagen, 79306) and purified with RNeasy mini kit (Qiagen, 74106) according to the kit protocol with DNase 1 treatment to exclude DNA contamination. Quality check was conducted with Agilent RNA 6000 Pico kit (cat no. 5067-1513) and mRNA-seq libraries were prepared as per the KAPA biosystem mRNA hyperprep (cat no. 08098123702). The sample was fragmented and the library quality was determined using the Agilent High Sensitivity D1000 kits (Cat no. 5067-5584 & 5067-5585). The library quantity was confirmed with KAPA library quantification kit protocol (KK4824) and the Illumina NovaSeq 6000 was used to sequence the samples with pair-end mode. After that, the sequences were mapped using Hisat2 to the mouse reference genome of ENSEMBL (version grcm38_snp_tran)(Kim, Langmead, & Salzberg, 2015). Differentially expressed genes were selected using DeSeq2 package (Love et al., 2014) and Metascape was used to perform Gene Ontology analysis(Z. Zhang et al., 2018).

## Funding and Acknowledgements

We are grateful to Professor Yichang Jia from the Tsinghua University, Beijing, for generously sharing embryonic thalamus samples of control and short TCF7L2 isoform knockout mice. Long TCF7L2 isoform knockout mouse model was generated at the facility headed by Prof. Andrzej Dziembowski at the University of Warsaw. Next-generation sequencing was performed at the Genomics Core Facility CeNT UW using the NovaSeq 6000 platform, financed by the Polish Ministry of Science and Higher Education (decision no. 6817/IA/SP/2018). We give special thanks to Krzysztof Goryca for the bioinformatic analysis of raw transcriptomic data. This work was supported by Narodowe Centrum Nauki (NCN, National Science Center) grants 2017/25/B/NZ3/01665 and 2020/37/B/NZ4/03261 to MBW. Additionally, AA and DP were supported by a NCN grant 2020/37/B/NZ2/03757 awarded to DP.

## Author contributions

MBW conceived and designed the study, analyzed and interpreted data, and drafted the manuscript. MOG designed the study, performed experiments, analyzed and interpreted data, and drafted the manuscript. JB performed ChIP-seq and contributed to data analysis and interpretation. MAL contributed to the design of the long Tcf7l2 KO model. AA performed bioinformatic analyses. EL contributed to data interpretation and drafted the manuscript. SB contributed to performing immunohistochemistry. HQ performed RNA-seq analysis and produced raw data. DP interpreted bioinformatic data. All authors critically read and approved the manuscript.

## Conflict of Interest

The authors declare no conflicts of interest, including conflicting financial interests, related to the work described

